# Structural and mechanistic insight into CRISPR-Cas9 inhibition by anti-CRISPR protein AcrIIC4

**DOI:** 10.1101/2021.07.09.451512

**Authors:** Sungwon Hwang, Chuxi Pan, Bianca Garcia, Alan R. Davidson, Trevor F. Moraes, Karen L. Maxwell

## Abstract

Phages, plasmids, and other mobile genetic elements express inhibitors of CRISPR-Cas immune systems, known as anti-CRISPR proteins, to protect themselves from targeted destruction. These anti-CRISPRs have been shown to function through very diverse mechanisms. In this work we investigate the activity of an anti-CRISPR isolated from a prophage in *Haemophilus parainfluenzae* that blocks CRISPR-Cas9 DNA cleavage activity. We determine the three-dimensional crystal struture of AcrIIC4 and show that it binds to the Cas9 Recognition Domain. This binding does not prevent the Cas9-anti-CRISPR complex from interacting with target DNA but does inhibit DNA cleavage. AcrIIC4 likely acts by blocking the conformational changes that allow the HNH and RuvC endonuclease domains to contact the DNA sites to be nicked.

## INTRODUCTION

CRISPR-Cas adaptive immune systems provide bacteria and archaea with protection from invasion by foreign genetic elements, including plasmids, transposons and phages.[1–3] These systems are widely distributed in nature, with approximately 40% of bacteria and 90% of archaea encoding at least one.[4,5] While CRISPR-Cas systems are extremely diverse, they all fall within one of two classes — Class 1 systems (types I, III, and IV) use multi-subunit surveillance complexes and Class 2 systems (types II, V, and VI) use single protein effectors to perform their function.[6] In this work we focus on the Class 2, type II-C system, which uses the Cas9 protein to carry out CRISPR-Cas activity.

Upon phage infection, CRISPR-Cas systems incorporate small fragments of the phage genome into the CRISPR array in a form known as a spacer.[7] The CRISPR array is transcribed and processed to form mature CRISPR RNA (crRNA) molecules, which bind to the Cas9 effector protein and serve as guides to target the complex to invading phage DNA for degradation in a sequence specific manner.[3] Once the phage DNA is recognized by the Cas9-crRNA complex, the target DNA strand is cleaved by an HNH endonuclease domain and the non-target strand is cleaved by a RuvC endonuclease domain.[8,9] The Cas9 protein accomplishes the various tasks required for CRISPR-Cas defense through the use of multiple functional domains. Cas9 can be broadly divided into two functional lobes; the α-helical recognition (REC) lobe and the nuclease (NUC) lobe. The nuclease lobe can be further functionally subdivided into the PAM-interacting domain (PID), and the HNH and RuvC endonuclease domains, which act to cleave the foreign DNA.

In response to the evolutionary pressures placed on them by CRISPR-Cas systems, phages have evolved anti-CRISPR proteins. Anti-CRISPR proteins allow phages to circumvent CRISPR-based immunity by inhibiting various components of the CRISPR-Cas system. The first anti-CRISPR proteins were discovered in phages infecting *Pseudomonas aeruginosa.*[10] Since then, over 90 distinct families of anti-CRISPR proteins have been identified and there are now anti-CRISPRs identified for every CRISPR-Cas type, including many for the Cas9-based type II systems that are widely used in genome editing.[11–14]

Anti-CRISPR proteins have been shown to obstruct Cas9 activity in a variety of different ways. For example, AcrIIC2 inhibits Cas9 binding to the CRISPR guide RNA, thereby blocking formation of the active ribonucleoprotein complex.[13] Anti-CRISPRs AcrIIA4 and AcrIIC5 block Cas9 target DNA binding,[15,16] while AcrIIC1 inhibits the HNH endonuclease domain and blocks the final DNA cleavage step.[12] In this study, we investigate the structure and function of anti-CRISPR AcrIIC4_*Hpa*_, which was identified in a *Haemophilus parainfluenzae* prophage[16]. We show that AcrIIC4_*Hpa*_ does not prevent target DNA binding but inhibits DNA cleavage, likely by sterically hindering the conformational changes that provide the endonuclease domains with access to the target DNA molecule.

## RESULTS

### AcrIIC4 binds to the REC2 domain

Previous work showed that AcrIIC4 blocks the activity of the Cas9 proteins from *H. parainfluenzae*, *Neisseria meningitidis, Simonsiella muelleri*, *Brackiella oedipodis* and *Kiloniella luminariae*.[16,17] We tested a second Cas9 variant from *N. meningitidis* (Nme2Cas9),[18] which shares 88% sequence identity with the previously examined *N. meningitidis* Cas9 (Nme1Cas9), for sensitivity to anti-CRISPR inhibition using an *in vivo* phage plaque assay. In this assay, Cas9 with an sgRNA targeting *Escherichia coli* phage Mu was co-expressed with AcrIIC4 in *E. coli.* In the absence of an anti-CRISPR, the targeting by Cas9-sgRNA leads to phage genome cleavage and prevents the phage from replicating. In the presence of an active anti-CRISPR, the Cas9 activity is inhibited and the ability of phage Mu to form plaques is restored. As can be seen in Figure 1(a), AcrIIC4 is able to robustly inhibit the activity of Nme2Cas9 to the same degree as Nme1Cas9.

**Figure 1.**
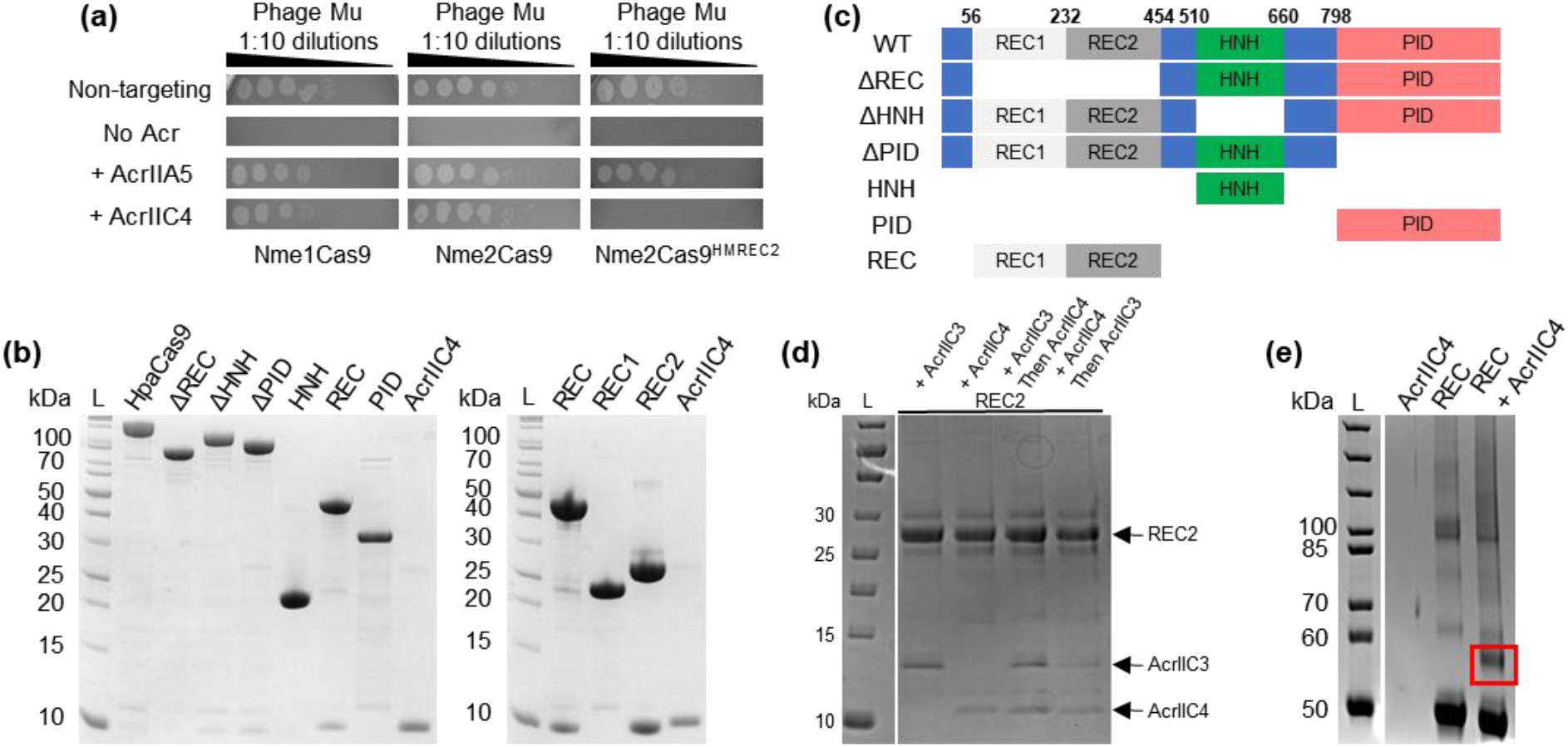
AcrIIC4 binds the REC2 domain. (a) Phage targeting assay with serial dilutions of phage Mu spotted onto lawns of *E. coli* expressing Nme1Cas9, Nme2Cas9, or Nme2Cas9 with the REC2 domain from *Neisseria* sp. HMSC070H10, sgRNA (non-targeting or targeting phage Mu), and anti-CRISPR (AcrIIA5 control or AcrIIC4). A representative figure from three replicates is shown. (b) N-terminal 6His-tagged HpaCas9 domains co-expressed with untagged AcrIIC4 were purified using nickel affinity chromatography. Purified proteins were analyzed by SDS-PAGE and visualized by Coomassie staining. Co-purification of untagged AcrIIC4 was detected by the presence of a ~10 kDa band. Purified AcrIIC4 is shown in the last lane of each gel. (c) Schematic of the HpaCas9 domain deletions and domain isolates used in the co-purification experiment shown in panel (b). (d) N-terminal 6His-tagged HpaCas9 REC2 was incubated with lysates of *E. coli* cells expressing untagged AcrIIC3 and AcrIIC4 and then purified by nickel affinity chromatography. Purified proteins were analyzed by SDS-PAGE and visualized by Coomassie staining. The proteins of interest are labeled. (e) Products of the cross-linking reaction were analyzed by SDS-PAGE and visualized by Coomassie staining. Control reactions of AcrIIC4 alone and HpaCas9 REC alone are shown in lane 2 and 3. The red box in lane 4 shows the band corresponding to crosslinked REC and AcrIIC4. The band was excised and analyzed by mass spectrometry.

To gain a better understanding of how AcrIIC4 inhibits these Cas9 proteins, we sought to determine the binding site of the anti-CRISPR protein. We first co-expressed untagged AcrIIC4 with 6His-tagged *H. parainfluenzae* (Hpa) Cas9 in *E. coli* and used nickel affinity chromatography to determine if the two proteins form a stable complex. When full-length 6His-HpaCas9 was purified, AcrIIC4 co-eluted with it from the Ni-NTA affinity column, demonstrating that the two proteins form a stable complex (Figure 1(b)). To determine which Cas9 domain AcrIIC4 binds to, we repeated the co-purification experiment using isolated Cas9 domains (PID, HNH and REC)(Figure 1(c)). No AcrIIC4 co-eluted with the PID and HNH domains, but a stable complex was observed when AcrIIC4 was co-expressed with the REC lobe (Figure 1(b)). Consistent with this, when the REC lobe was deleted from the full-length Cas9 protein, AcrIIC4 was no longer able to bind (Figure 1(b)). In type II-C Cas9 proteins, the REC lobe is comprised of two domains, REC1 and REC2.[19] By co-expressing the isolated REC1 and REC2 domains, we found that AcrIIC4 co-eluted with REC2, but not REC1 (Figure 1(b)). This showed that the REC2 domain alone is sufficient for AcrIIC4 binding.

To provide further evidence that the REC2 domain is the site of interaction, we engineered a chimeric version of Nme2Cas9 where the native REC2 domain was replaced by the REC2 domain of the Cas9 from *Neisseria* sp. HMSC070H10, which shares 88% sequence identity with Nme2Cas9 overall, but only 38% sequence identity in the REC2 domain. This chimeric Cas9 (Nme2Cas9^HMREC2^) was able to inhibit replication of phage Mu in the phage-targeting assay, and the expression of AcrIIC4 did not relieve this inhibition (Figure 1(a)). Expression of the broad-specificity AcrIIA5, which induces truncation of the sgRNA,[17] restored the ability of phage Mu to form plaques, showing that this chimeric Cas9 can be inhibited by anti-CRISPR activity (Figure 1(a)). These results provide additional evidence that the REC2 domain is the primary binding site for AcrIIC4.

Previously, anti-CRISPR protein AcrIIC3 was also shown to bind to the REC2 domain.[19,20] To determine if AcrIIC3 and AcrIIC4 compete for a common binding site on the REC2 domain or if they can bind simultaneously, we conducted another nickel affinity co-purification experiment with purified HpaCas9 REC2 and lysates containing untagged AcrIIC3 and AcrIIC4. We found that AcrIIC3 and AcrIIC4 could bind simultaneously, as shown by the co-elution of both proteins when incubated with the REC2 domain (Figure 1(d)). The amount of each anti-CRISPR binding to the REC2 domain was equal to that observed when they were incubated with the REC2 domain individually, and the order of addition did not change their binding profiles (Figure 1(d)). These results demonstrate that AcrIIC3 and AcrIIC4 bind to different regions of the REC2 domain.

In an attempt to further elucidate the precise binding site of AcrIIC4, we performed a protein cross-linking experiment with the HpaCas9 REC lobe and AcrIIC4. We purified the REC lobe and AcrIIC4 separately, mixed the purified proteins together and added disuccinimidyl suberate (DSS), a chemical crosslinker that is reactive to primary amines (i.e. N-termini or lysine side chains).[21] The cross-linked reaction products were visualized by SDS-polyacrylamide gel electrophoresis followed by Coomassie staining (Figure 1(e)). A band with a size corresponding to that predicted to result from the crosslinking of the REC lobe and AcrIIC4 in a 1:1 ratio was excised from the gel and subjected to trypsin digestion. The resulting peptides were analyzed by tandem mass spectrometry (MS/MS) to locate the crosslinks. MS/MS analysis revealed five unique crosslinks between the REC lobe and AcrIIC4. These corresponded to two residues in the REC lobe (Lys78 and Lys135) cross linking with residues in the anti-CRISPR (Lys78-Lys25, Lys78-Lys61, Lys135-Lys25, Lys135-Lys50, Lys135-Lys61). Both residues on the REC lobe, Lys78 and Lys135, are found in the REC1 domain. As our previous data showed that the REC2 domain alone is sufficient for AcrIIC4 binding, these crosslinking data suggest that the precise binding site is likely located between the REC1 and REC2 domains.

### AcrIIC4 inhibits DNA cleavage but allows DNA binding

To investigate the effects of AcrIIC4 activity on DNA binding, we set up an *in vivo* transcriptional repression assay. This assay uses a constitutively expressed artificial promoter that drives expression of the lux operon in *E. coli*.[17] When Nme1Cas9 is co-expressed with an sgRNA that targets the lux operon promoter and is not catalytically inhibited by an anti-CRISPR, it cleaves the promoter leading to cell death. (Figure 2(a, top)). When co-expressed with an anti-CRISPR that does not block DNA binding but prevents DNA cleavage, Nme1Cas9 binds to the promoter region and blocks transcription of the lux operon, leading to low levels of luminescence (Figure 2(a, middle)). By contrast, co-expression of an anti-CRISPR that prevents DNA binding relieves repression of the lux operon and leads to increased luminescence (Figure 2(a, bottom)).

**Figure 2.**
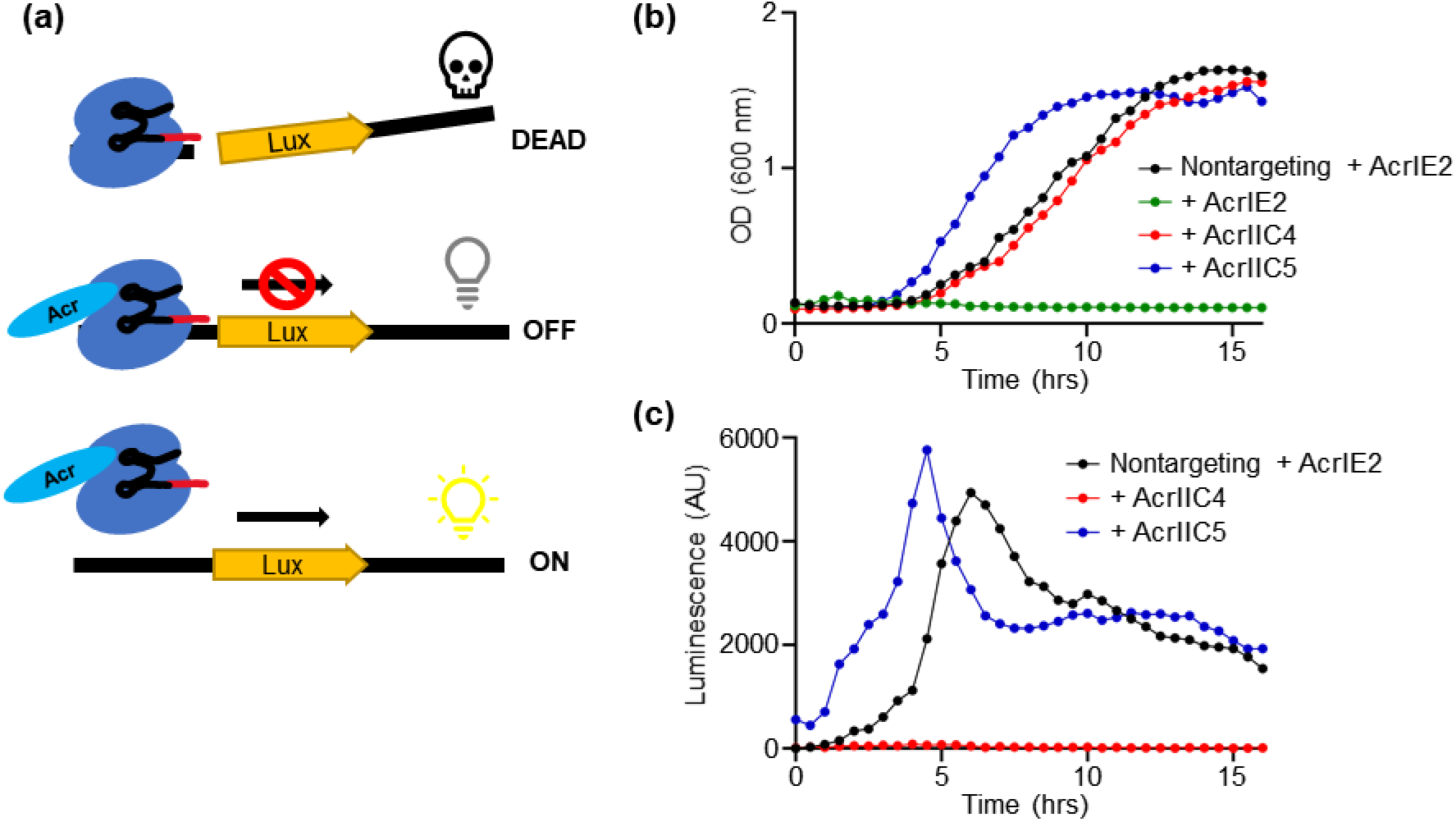
Luminescence transcriptional repression assay. (a) Schematic of the assay. Without inhibition by an anti-CRISPR protein, Nme1Cas9 will target and cleave the plasmid containing the lux operon, resulting in cell death. When inhibited by an anti-CRISPR that does not block DNA binding, Nme1Cas9 will bind to the lux operon promoter and repress transcription, blocking luminescence. When inhibited by an anti-CRISPR that blocks DNA binding, Nme1Cas9 will not bind to the promotor, resulting in transcription of the lux operon and luminescence. (b) Growth curves of cells where Nme1Cas9 was co-expressed with non-targeting sgRNA (black) or inhibited by AcrIIC4 (red) or AcrIIC5 (blue). AcrIE2 does not inhibit Nme1Cas9, resulting in cell death (green). A representative plot of at least three replicates is shown. (c) Graphs of luminescence signal over time. When Nme1Cas9 was co-expressed with non-targeting sgRNA (black) or AcrIIC5 (blue) that blocks DNA binding, luminescence signal was detected. No luminescence was seen when Nme1Cas9 was co-expressed with AcrIIC4 (red), indicating that AcrIIC4 is not blocking DNA binding.

When Nme1Cas9 was co-expressed with AcrIIC4, AcrIIC5_*Smu*_, or an sgRNA that does not target the lux operon promoter, the *E. coli* cells were able to grow normally, as expected, due to inhibition of Cas9 catalytic activity (Figure 2(b)). By contrast, when Nme1Cas9 was co-expressed with targeting sgRNA and AcrIE2, an anti-CRISPR that does not inhibit Cas9,[10,22] the cells died as a result of the Nme1Cas9 activity (Figure 2(b)). For the cells that grew normally, luminescence was detected when Nme1Cas9 was co-expressed with non-targeting sgRNA or with targeting sgRNA and AcrIIC5_*Smu*_, which was previously shown to block target DNA binding (Figure 2(c)).[16] By contrast, luminescence was not detected when Nme1Cas9 was co-expressed with targeting sgRNA and AcrIIC4 (Figure 2(c)). These results show that AcrIIC4 inhibits the cleavage activity of Nme1Cas9 as the cells are able to survive. They further show that AcrIIC4 allows binding of Nme1Cas9 to target DNA as the complex repressed luminescence in this assay.

To further probe the DNA-binding activity of Nme1Cas9 in the presence of AcrIIC4 we assessed DNA binding *in vitro* through the use of electrophoretic mobility shift assays. We first co-expressed Nme1Cas9 and sgRNA in *E. coli* and purified the ribonucleoprotein complex. The purified complex was then incubated with increasing amounts of purified AcrIIC4, fluorescein labeled target DNA was added, and an electrophoretic mobility shift assay was performed. The binding of the Cas9-sgRNA complex to a 44 bp dsDNA fragment containing a protospacer sequence complementary to the sgRNA spacer caused a change in mobility of the dsDNA fragment in a polyacrylamide gel (Figure 3(a)). Addition of AcrIIC4 to the DNA-bound Cas9 complex did not result in a decrease in DNA binding at up to 50-fold excess of AcrIIC4 (Figure 3(a)). By contrast, AcrIIC5, which inhibits DNA binding,[16] showed decreased target DNA binding with increasing anti-CRISPR concentration (Figure 3(b)). Although the Nme1Cas9-AcrIIC4 complex was able to bind to target DNA, this complex did not mediate DNA cleavage *in vitro* (Figure 3(c)).

**Figure 3.**
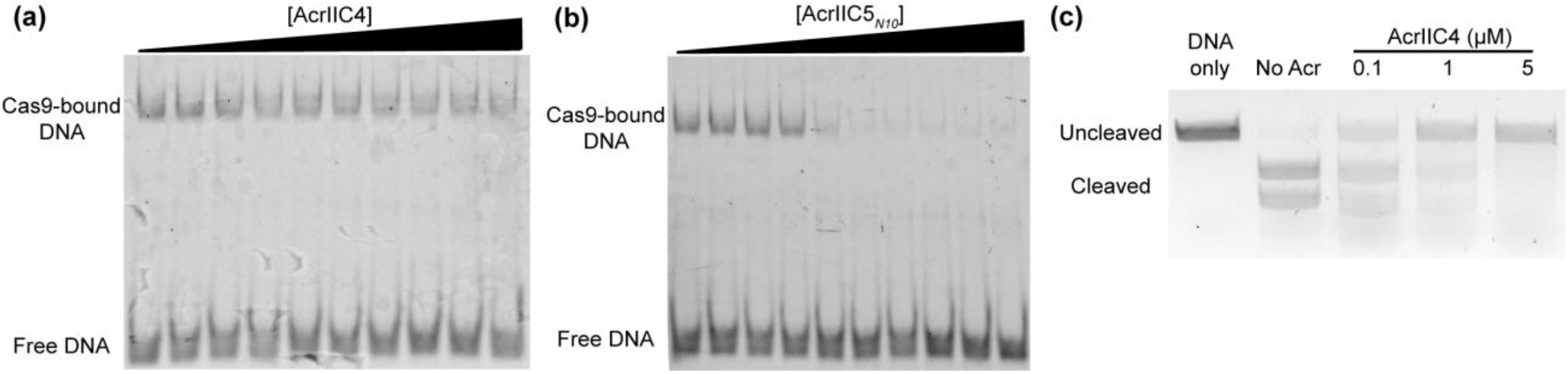
AcrIIC4 does not inhibit target DNA binding. (a) Electrophoretic mobility shift assay (EMSA) with 1 μM Nme1Cas9-sgRNA RNP, 100 nM fluorescein labelled target DNA, and increasing AcrIIC4 concentrations. The assay was analyzed by native PAGE, and the DNA was visualized using a Chemidoc imager. The amount of Cas9-bound DNA was unchanged even at the highest concentrations of AcrIIC4. (b) EMSA with 1 μM Nme1Cas9-sgRNA RNP, 100 nM fluorescein labelled target DNA, and increasing AcrIIC5_*N10*_ concentrations. Unlike with AcrIIC4, the amount of Cas9-bound DNA decreases with increasing concentrations of AcrIIC5_*N10*_. (c) DNA cleavage assay with 150 nM Nme1Cas9-sgRNA RNP, 300 ng of target plasmid DNA, and AcrIIC4 at 0, 0.1, 1, or 5 μM. The reactions were analyzed on a 1 % agarose gel supplemented with SYBR Safe. With increasing AcrIIC4 concentration, more target DNA is left uncleaved.

While we showed that AcrIIC4 interacts directly with Cas9 through the REC2 domain (Figure 1), we did not know if it remained bound to Cas9 following DNA binding. Previous studies have shown that some anti-CRISPR proteins act via enzymatic means and do not remain stably bound to the Cas9 complex.[23–25] To determine if AcrIIC4 remained bound in the presence of DNA, we mixed AcrIIC4-bound Nme1Cas9-sgRNA with a 44 base pair target DNA oligomer and used size-exclusion chromatography to analyze the equilibrium complex (Figure 4(a)). SDS-polyacrylamide gel electrophoresis followed by Coomassie staining was used to visualize the protein components (Figure 4(b)) and urea-PAGE and SYBR gold staining for the sgRNA and DNA (Figure 4(c)). We observed three peaks eluting from the size exclusion column. The first peak contained Nme1Cas9 and AcrIIC4 (Figure 4(b)) as well as sgRNA and DNA (Figure 4(c)). The second peak contained free DNA (Figure 4(c)), and the third peak contained excess AcrIIC4 (Figure 4(b)). These results provide additional evidence that AcrIIC4 does not affect DNA binding by Nme1Cas9 and suggest that all four components form a stable complex.

**Figure 4.**
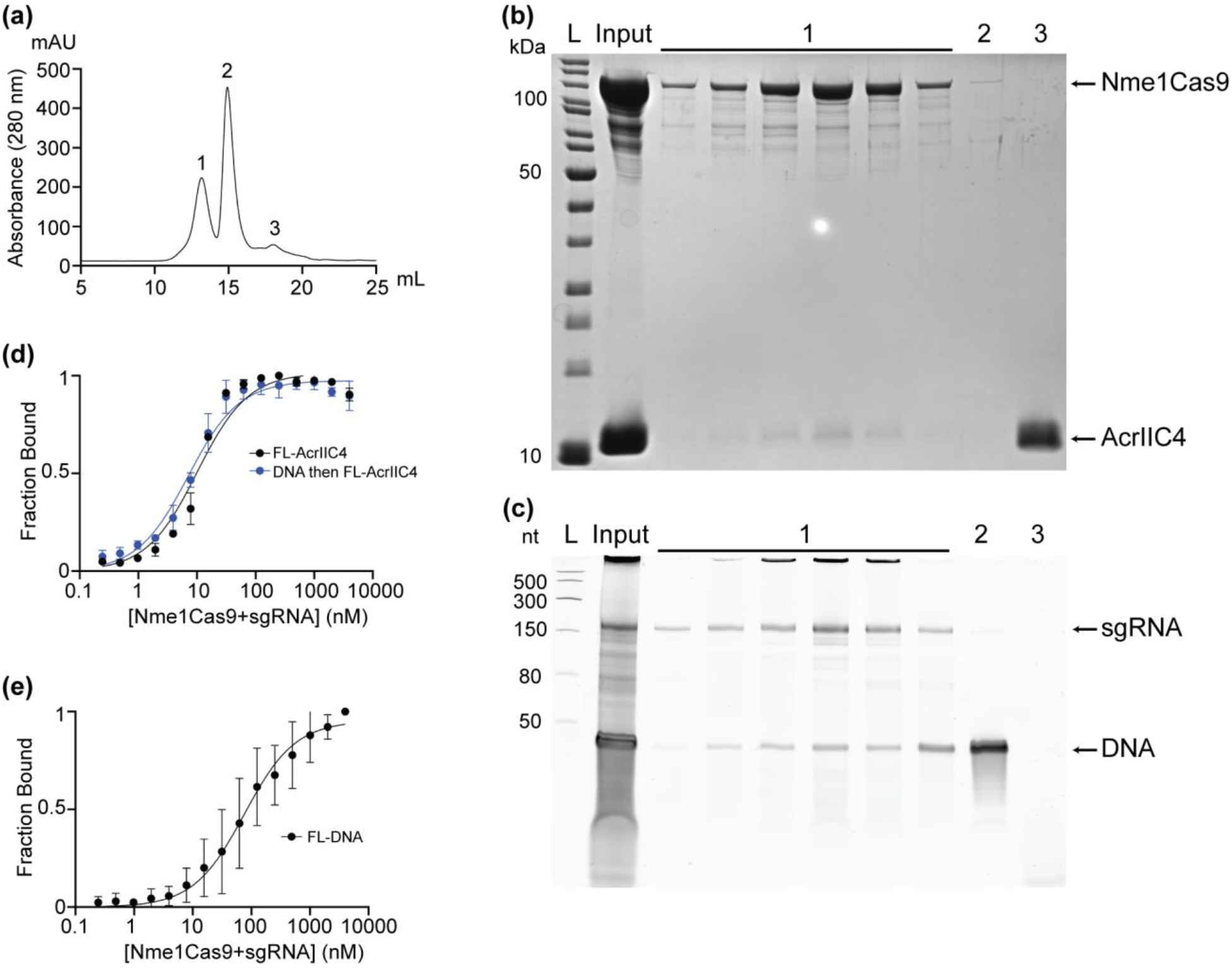
AcrIIC4 can bind to Cas9 simultaneously with target DNA. (a) A_280 nm_ chromatogram of the Nme1Cas9-sgRNA-DNA-AcrIIC4 complex run on a Superdex 200 Increase 10/300 GL gel filtration column. A total of three peaks appeared in the chromatogram. (b) SDS-PAGE gel stained with Coomassie of the chromatogram peaks. In peak 1, Nme1Cas9 and AcrIIC4 co-eluted. In peak 2, there is no significant protein present. Excess AcrIIC4 eluted in peak 3. (c) Urea-PAGE gel stained with SYBR Gold of the chromatogram peaks. In peak 1, both sgRNA and DNA co-eluted with the complex. Excess DNA is present in peak 2. There is no nucleic acid present in peak 3. (d) Fluorescence polarization (FP) binding curves of fluorescein (FL) labelled AcrIIC4 and Nme1Cas9-sgRNA RNP with (blue) and without (black) pre-incubation with target DNA. (e) FP binding curve of FL labelled target DNA with Nme1Cas9 RNP. All FP curves show results from three replicates with the standard deviation plotted as error bars.

To determine if there were differences in the binding affinity of AcrIIC4 to Cas9 in the presence and absence of bound dsDNA we used a fluorescence polarization assay. In this assay the binding of fluorescently labeled AcrIIC4 to the Cas9-sgRNA complex reduces its tumbling rate and leads to an increase in the fluorescence polarization signal. This allows us to directly compare the binding of AcrIIC4 to Cas9-sgRNA in the presence and absence of dsDNA. Consistent with the EMSA results, the binding of AcrIIC4 to both Cas9-sgRNA alone and in complex with dsDNA was readily detectable (Figure 4(d)). The equilibrium binding constants (K_*d*_) were determined for the binding of fluorescently-labelled AcrIIC4 to Nme1Cas9-sgRNA complex alone and bound to target DNA (Figure 4(d)). We found no significant difference in the calculated K_*d*_ values (10.2 ± 2.0 nM alone vs. 7.1 ± 1.3 nM with DNA), showing that AcrIIC4 binds equally well to the Nme1Cas9-sgRNA complex in its apo and DNA bound states. A control assay adding increasing amounts of Nme1Cas9-sgRNA to fluorescently-labelled target DNA confirmed that a stable interaction forms between Cas9-sgRNA and DNA in these assay conditions (Figure 4(e)). The calculated K_*d*_ for this experiment was 75.9 ± 38.9 nM, which is similar to previously reported values.[12,16] Combined, these results show that AcrIIC4 does not block DNA binding, and thus must inhibit Cas9 at the DNA cleavage step.

### The crystal structure of AcrIIC4

To better understand the mechanism of AcrIIC4 activity, we determined its crystal structure to a resolution of 1.73 Å using iodide—single-wavelength anomalous diffraction (I-SAD). The X-ray data collection and refinement statistics are presented in Table 1. The 88-residue AcrIIC4 protein crystallized as a homodimer, with one helix from each monomer contributing to a single long coiled interface (Figure 5(a-c)). The interaction of these helices forms an extended hydrophobic interaction interface composed of 23 hydrophobic residues with side chains that are greater than 85% buried. The monomers are packed in a head to tail fashion, with two short helical segments (residues Ser5-Ile11 and Ser14-Val22) located at the N-termini of each protein. Residues within these two helical segments pack against the long extended helices of each monomer (residues Ly25-Gln84), forming a hydrophobic core with residues contributed by the N-terminal region of the one monomer and the C-terminal region of the other monomer (Figure 5(d)). This results in the formation of a globular domain at each end of the long, extended helical bundle, an arrangement that resembles a dumbbell-like structure. In addition to these domains, hydrophobic residues from each monomer pack together along the length of the extended helices and contribute additional stability to the dimeric interface (Figure 5(d)). A search of the structural database using DALI[26] revealed no similarities between AcrIIC4 and any previously determined protein structure that might suggest an evolutionary or functional relationship.

**Table 1.**
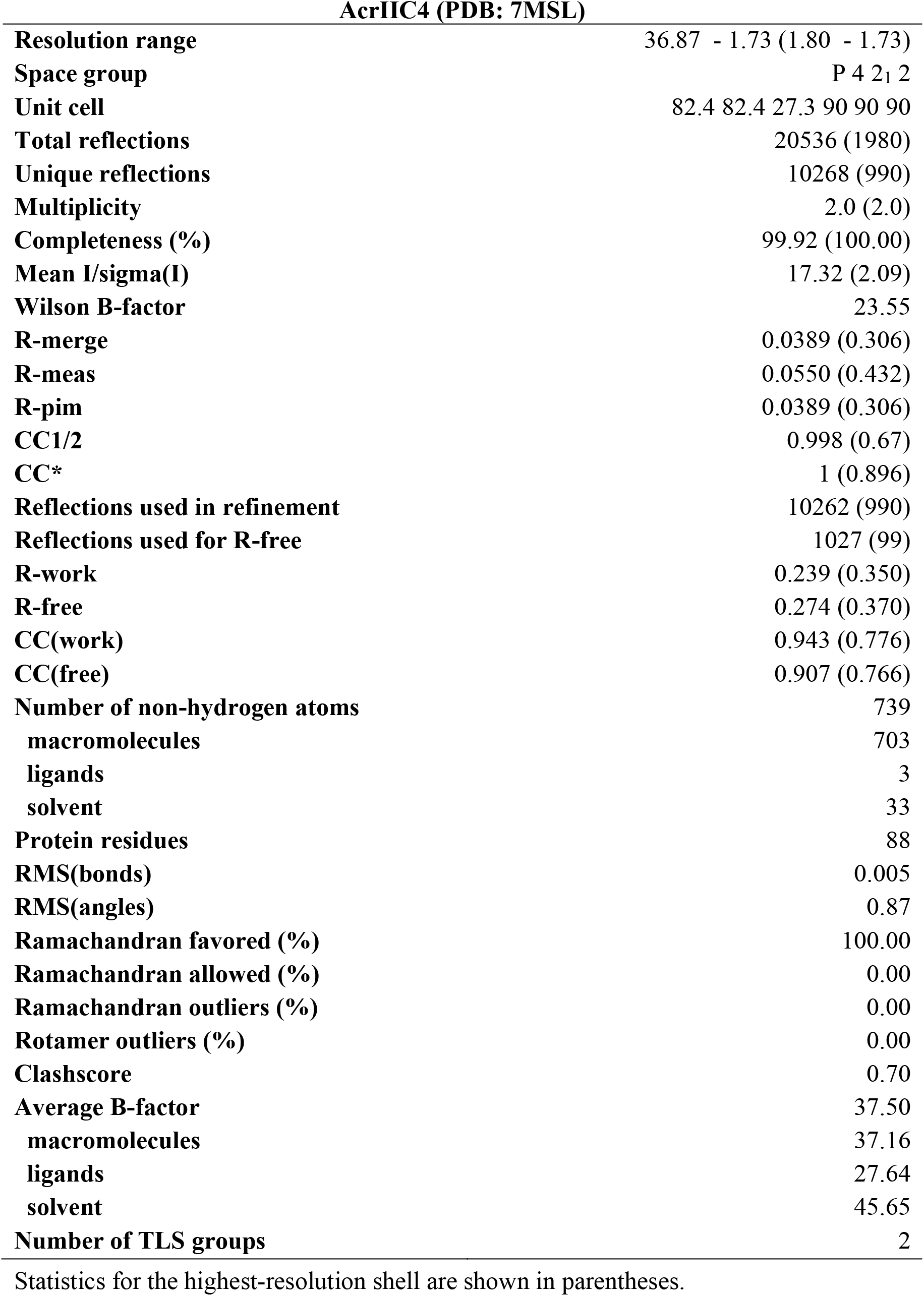
Data collection and refinement statistics.

**Figure 5.**
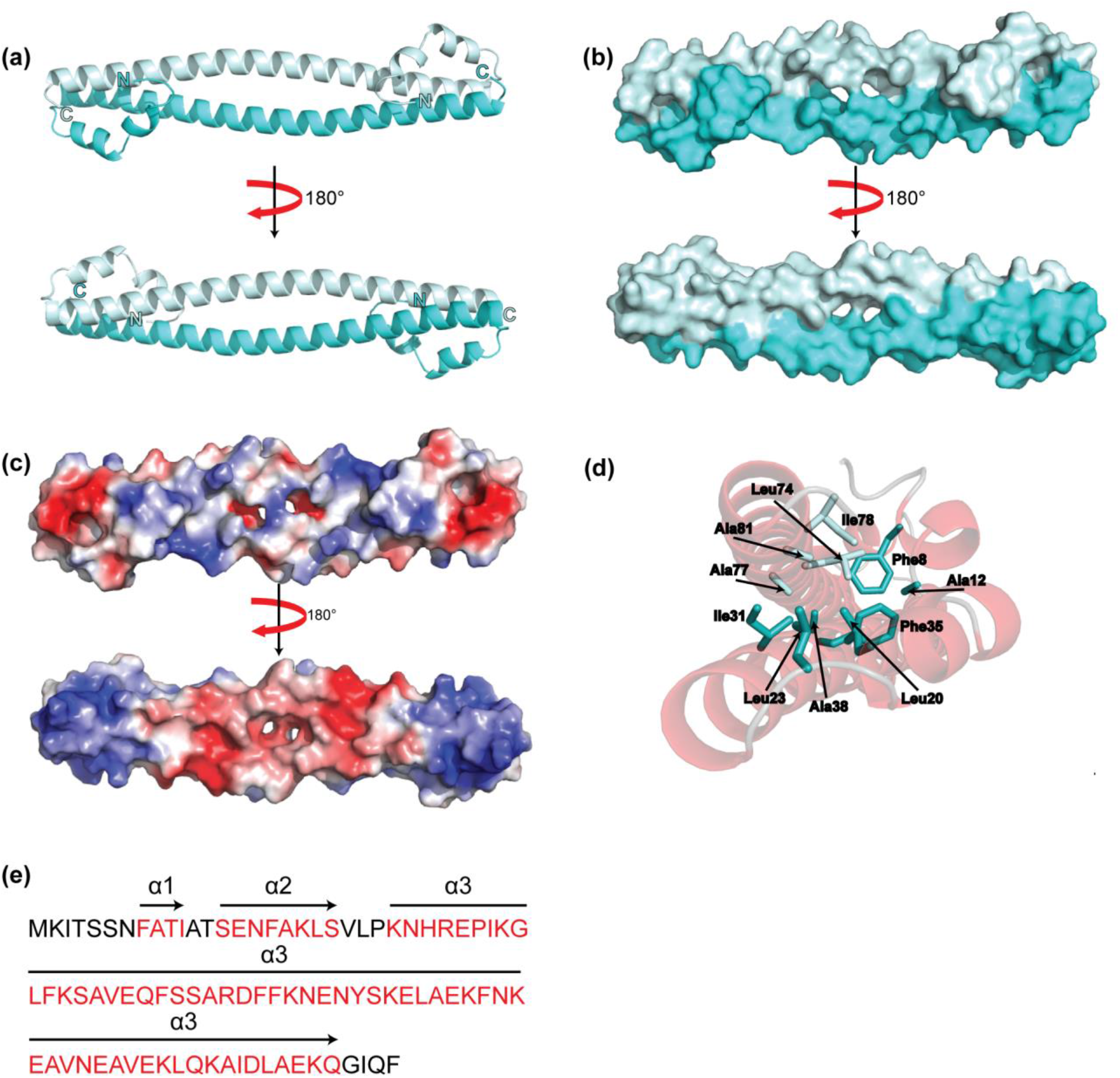
Crystal structure of AcrIIC4. (a) The structure of AcrIIC4 with cartoon representation. The light blue and cyan show each monomer with the N- and C-termini labelled. (b) Surface representation of AcrIIC4. (c) Surface representation of AcrIIC4 displaying the electrostatics ranging from red (electronegative) to blue (electropositive), generated using the Adaptive Poisson-Boltzmann Solver (APBS) plugin in PyMOL. (d) The hydrophobic pocket of AcrIIC4. The side chains of residues from both monomers (light blue and cyan) that contribute to the hydrophobic core are labelled. (e) AcrIIC4 amino acid sequence. The red-colored amino acids and the arrows above represent the three alpha helices that make up the structure.

### Identification of a functionally important surface of AcrIIC4

The lack of sequence homologues of AcrIIC4 made it difficult to predict residues that might be important for mediating the interaction with Cas9. Thus, we used the structure to identify 18 solvent-exposed residues distributed across the surface of the protein and substituted an alternative amino acid at each position (Table 2, Figure 6(a)). The inhibitory activities of these AcrIIC4 mutants were then tested using the *in vivo* phage targeting assay. We found that three of the 18 substitutions on the surface of AcrIIC4 led to a decrease in anti-CRISPR activity greater than 100-fold (Table 2). These residues (Arg46, Phe49, and Lys56) cluster on one side of the AcrIIC4 dimer (Figure 6(a)), suggesting that this may be an important functional surface of the protein.

**Table 2.** Nme1Cas9 phage targeting assay with AcrIIC4 mutants.

| Target Residue | Mutation | Fold Reduction Compared to WT |
| --- | --- | --- |
|  |  | Nme1Cas9 |
| K19 | K19A | $10^2$ |
| K25 | K25A | 0 |
| K36 | K36D | $10^2$ |
| E40 | E40T | 10 |
| R46 | R46A | $10^5$ |
| F48 | F48A | 0 |
| F49 | F49A | $10^8$ |
| K50 | K50A | 0 |
|  | K50D | 0 |
| Y54 | Y54A | 10 |
| K56 | K56A | $10^4$ |
| K61 | K61A | 0 |
|  | K61E | 0 |
| K64 | K64E | 0 |
| E65 | E65A | 0 |
|  | E65T | 10 |
| E69 | E69T | 0 |
| E72 | E72K | 10 |
| K76 | K76A | 0 |
| D79 | D79T | 0 |
| E82 | E82K | 0 |

**Figure 6.**
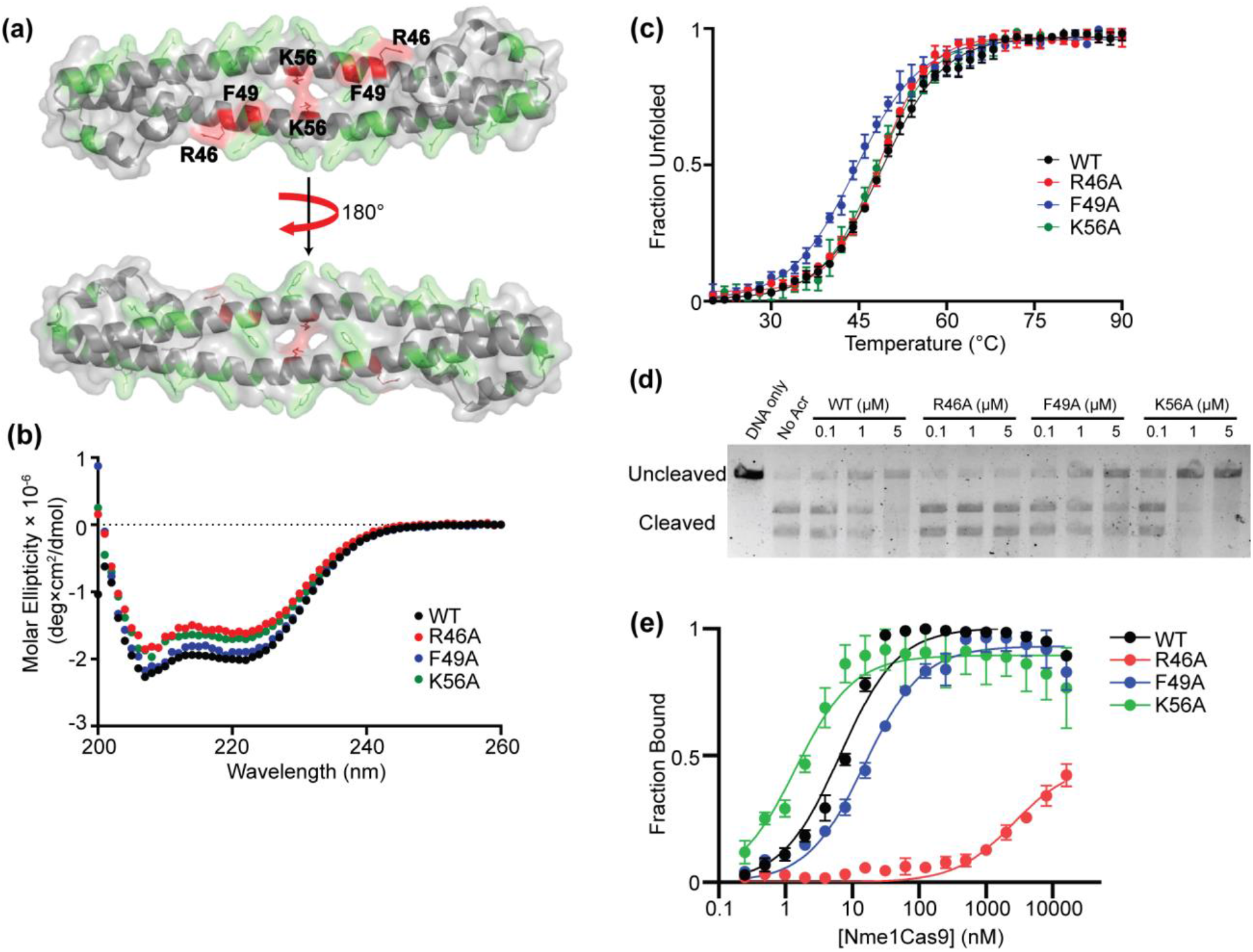
The functional surface of AcrIIC4. (a) Surface representation of the AcrIIC4 structure with the functionally important residues labelled. The amino acids that were found to have minimal or no effect on AcrIIC4 activity are colored in green. Arg46, Phe49, and Lys56 (red) are all located on a common surface of AcrIIC4 and make up a potential functionally important region. (b) Far-UV circular dichroism spectra of AcrIIC4 mutants. A representative plot of at least three replicates is shown. The mutants’ spectrum profiles are similar to the wild type’s, indicating that the mutations do not disturb the structural integrity of the protein. (c) Temperature-induced unfolding curves of AcrIIC4 mutants. The melting temperatures (T_m_) of the R46A (red, 48.5 ± 0.3 °C), F49A (blue, 44.4 ± 0.5 °C), and K56A (green, 48.1 ± 0.4 °C) mutants were all within 3 °C of wild type (49.0 ± 0.3 °C). The plot show results from three replicates with the standard deviation plotted as error bars. (d) DNA cleavage assay with 150 nM Nme1Cas9-sgRNA RNP, 300 ng of target plasmid DNA, and wild type or mutant AcrIIC4 at 0, 0.1, 1, or 5 μM. The reactions were analyzed on a 1 % agarose gel supplemented with SYBR Safe. There is a significant reduction of Cas9 nuclease activity inhibition by the R46A variant, and partial reduction by the F49A variant. A representative figure from three replicates is shown. (e) Fluorescence polarization binding curves of fluorescein labelled wild type and mutant AcrIIC4 with Nme1Cas9. Only the R46A substitution caused a significant reduction of affinity compared to wild type AcrIIC4. The plot shows results from three replicates with the standard deviation plotted as error bars.

To ensure that the decrease in anti-CRISPR activity we observed was due to a weakened protein-protein interaction and not due to misfolding or decreased stability, we used circular dichroism spectroscopy to assess the thermodynamic stabilities of the variant proteins. We expressed 6-His tagged versions of the Arg46, Phe49 and Lys56 variants in *E. coli* and purified them using Ni-affinity chromatography. As shown in Figure 6(b), the far-UV circular dichroism spectra were very similar to the spectrum of the wild type AcrIIC4 protein, indicating that there were no large changes in the secondary structure of these proteins as compared with wild type. We also examined the thermal stability of these mutants using temperature-induced unfolding experiments monitored by CD and discovered that the melting temperatures of these mutants were within 3°C of wild type AcrIIC4 (49.0 ± 0.3°C) (Figure 6(c)). These data show that the mutants with decreased activity maintain a well folded tertiary structure.

We next examined whether the AcrIIC4 mutants inhibited Nme1Cas9 DNA cleavage activity *in vitro*. Nme1Cas9-sgRNA was incubated with wild-type or mutant AcrIIC4 and then a linearized plasmid containing the protospacer and PAM sequence targeted by the sgRNA was added. In the absence of AcrIIC4, the DNA fragment was cleaved in a Cas9-sgRNA dependent manner (Figure 6(d)). The addition of wild type AcrIIC4 inhibited this cleavage activity. We observed a loss of anti-CRISPR activity with the R46A variant, and a partial loss of activity with the F49A variant. By contrast, the K56A variant inhibited Nme1Cas9 nuclease activity at levels similar to wild-type AcrIIC4. We next examined the effects of these mutations on binding to apo-Cas9 using the fluorescence polarization assay. The *K*_d_ of wild-type AcrIIC4 binding to Nme1Cas9 was determined to be 6.6 ± 1.1 nM (Figure 6(e)). The R46A substitution caused a ~400-fold decrease in affinity (>2.5 μM). There was only a ~2-fold decrease in affinity for the F49A substitution (15.3 ± 2.6 nM), while there was actually a slight increase in affinity for the K56A substitution (1.4 ± 0.4 nM). The reasons for the disparity between this assay, which shows similar binding affinities for wild type AcrIIC4 and the F49A and K56A mutants, and the *in vivo* assay, which shows decreased activity of these mutants, are not clear. Taken together, these results suggest an important role for Arg46 and show that the ability to stably bind the Cas9-sgRNA complex is required for anti-CRISPR activity.

As our cross-linking data suggested that AcrIIC4 binds between the REC1 and REC2 domains, we examined the Nme1Cas9-sgRNA structure (PDB: 6JDQ)[19] and identified a highly electronegative pocket on a region of the REC2 domain adjacent to the REC1 domain (Figure 7(a)). We hypothesized that this electronegative surface might bind the positively charged region of AcrIIC4 surrounding the R46 position. To test this, we individually substituted four negatively charged residues, Asp392, Glu393, Asp394, and Glu418, with either alanine or lysine and tested the ability of AcrIIC4 to inhibit these mutant Nme1Cas9 proteins using the *in vivo* phage targeting assay. We found that AcrIIC4 activity was reduced by >1000-fold against the Cas9^D394K^ variant while the other variants were unaffected (Figure 7(b)). Cas9^D394K^ was able to robustly inhibit phage plaquing in the absence of AcrIIC4, showing that this mutation did not affect Cas9 activity. Furthermore, Cas9^D394K^ was inhibited by AcrIIC1, which interacts with the HNH domain to prevent DNA cleavage, demonstrating that the reduction of Cas9-inhibition is specific to AcrIIC4 (Figure 7(b)). These results suggest that Asp394 forms part of the interaction surface to which AcrIIC4 binds.

**Figure 7.**
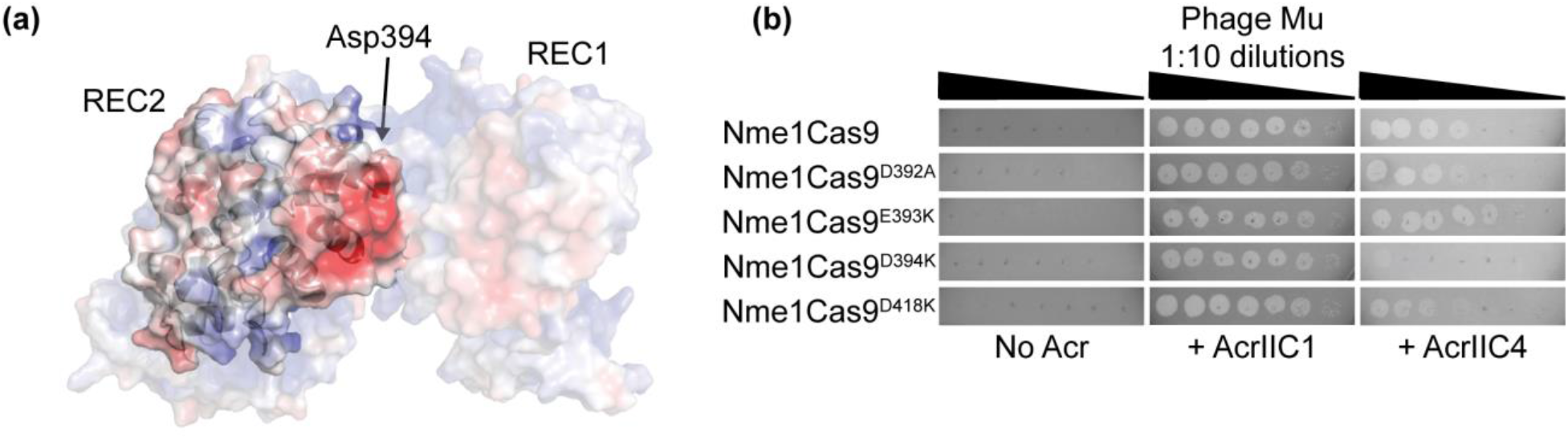
AcrIIC4 activity is greatly reduced against Nme1Cas9^D394K^ variant. (a) Surface representation of the Nme1Cas9 structure in complex with sgRNA (PDB: 6JDQ) displaying the electrostatics ranging from red (electronegative) to blue (electropositive), generated using the Adaptive Poisson-Boltzmann Solver (APBS) plugin in PyMOL. The figure highlights the highly electronegative pocket on a region of the REC2 domain adjacent to the REC1 domain. (b) Phage targeting assay with serial dilutions of phage Mu spotted onto lawns of *E. coli* (gray background) expressing wild type Nme1Cas9 or Nme1Cas9 variant, sgRNA targeting phage Mu, and anti-CRISPR (AcrIIC1 control or AcrIIC4). The lighter gray circles indicate zones of clearing of bacteria caused by successful infection and propagation of phage Mu. All Nme1Cas9 variants inhibit phage Mu propagation without anti-CRISPR, demonstrating that the point mutations do not affect Cas9 activity. The reduced clearing by phage Mu with AcrIIC4 indicates reduction of inhibitory activity against the Cas9^D394K^ variant. A representative figure of three replicates is shown.

### AcrIIC4 may inhibit important conformational changes in Cas9

The ability of AcrIIC4 interact with both free and DNA-bound Cas9-sgRNA through the REC lobe suggested that it may inhibit DNA cleavage by an allosteric mechanism. Cas9 is known to undergo a number of conformational changes that are required for DNA cleavage.[19,27] In the final stages of CRISPR-Cas activity, Cas9 binds to the target DNA through complementary base pairing with the guide RNA and the HNH and RuvC endonuclease domains can then access and cleave the target DNA. A previous crystallographic study of Nme1Cas9 showed that the HNH endonuclease domain must undergo a large conformational shift to access the DNA cleavage site. This is accompanied by a simultaneous outward rotation of the REC2 domain to make room for the HNH domain.[19] We hypothesize that AcrIIC4 binding to the REC2 domain prevents this motion, thus locking the Cas9 in a catalytically inactive state.

To investigate this hypothesis, we modelled our AcrIIC4 structure onto the previously determined Nme1Cas9 structures[19] using the HADDOCK (High Ambiguity Driven protein-protein DOCKing) web server.[28] This program creates models of biomolecular complexes using a flexible docking approach that takes into account information about experimentally determined protein interfaces. We began with the Nme1Cas9 structure in complex with sgRNA and target DNA in a seed base pairing state (PDB: 6KC7). In this state the sgRNA is partially duplexed with target DNA; guide nucleotides 1-16 and parts of the REC2 domain are disordered and the HNH domain active site is not located near the target DNA. We designated Arg46 from AcrIIC4 and Asp394 from Nme1Cas9 as protein interface residues and used HADDOCK to generate a set of low energy Nme1Cas9-AcrIIC4 complex structures. These structures could be broadly categorized into five clusters based on their conformation (Supplementary Figure 1). We then closely examined these groups of structures to determine which ones were consistent with the cross-linking data showing the REC lobe residues Lys78 and Lys135 were in close proximity to AcrIIC4 residues Lys25, Lys50, and Lys61. One cluster of structures had these residues in close proximity, with an average distance of ~25 Å, which is within the 26-30 Å distance constraint of DSS crosslinking.[29] These structures were selected for further analysis.

In this model, AcrIIC4 is nestled into a crevice between the REC1 and REC2 subdomains (Figure 8(a)). In this position, AcrIIC4 does not interfere with the known binding interface between REC2 and AcrIIC3, which is consistent with our experimental results that showed AcrIIC3 and AcrIIC4 can bind the REC lobe simultaneously (Supplementary Figure 2). We modeled this conformation of AcrIIC4 into the Nme1Cas9 structure in the catalytically active state (PDB: 6JDV), keeping the anti-CRISPR (cyan) and aligning the Nme1Cas9 structures (Figure 8(b)). In the catalytically active state, the REC2 (black) and HNH (green) domains show large conformational changes, moving inward towards the DNA. This movement leads to the formation of a number of steric clashes between AcrIIC4 and Nme1Cas9, particularly in the REC2 and HNH domains (Figure 8(c); red hexagons), that are not present in the original catalytically active structure (Figure 8(d); red hexagons). These steric clashes resulting from the presence of AcrIIC4 bound to the REC domain would prevent the HNH endonuclease domain from reaching the target DNA, locking the Cas9 in the inactive DNA-bound state. This model supports our hypothesis that AcrIIC4 may inhibit Cas9 activity by preventing conformational changes that are required to cleave the foreign DNA molecule.

**Figure 8.**
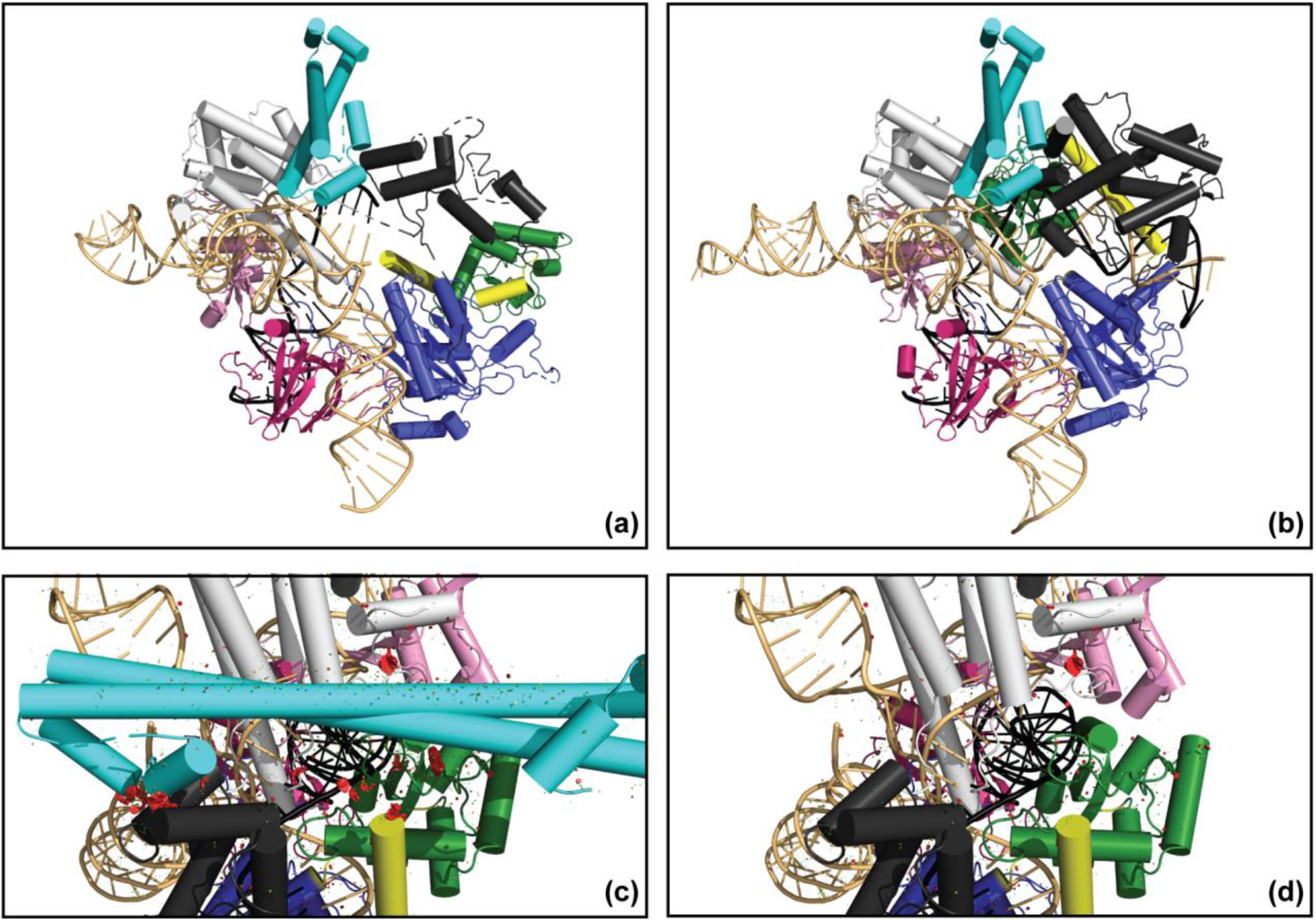
AcrIIC4 may inhibit important conformational changes in Cas9. (a) Model of AcrIIC4 (cyan) on the Nme1Cas9 structure in complex with sgRNA (tan) and target DNA (black) in a seed base pairing state (PDB: 6KC7) generated using the HADDOCK web server. Nme1Cas9 is colored by domain (RuvC – blue, REC1 – white, REC2 – black, linkers – yellow, HNH – green, WED – pink, PID – dark pink). AcrIIC4 is modelled in a crevice between the REC1 and REC2 domains. (b) Model of AcrIIC4 projected onto the Nme1Cas9 structure in the catalytically active state (PDB: 6JDV). (c) Close-up of (b) with steric clashes between the modelled AcrIIC4 (cyan) and the REC2 (black) and HNH (green) domains of Nme1Cas9 represented with red hexagons. The size of the hexagons is proportional to the amount of steric strain. (d) Close-up of the Nme1Cas9 structure in the catalytically active state (PDB: 6JDV). The steric clashes are not observed in the absence of AcrIIC4.

## DISCUSSION

In this work we investigated the mechanism of activity of anti-CRISPR protein AcrIIC4 and show that it blocks Cas9 activity after DNA binding. We found that the REC2 domain was the primary site of interaction, and that both sgRNA loading and dsDNA binding occurred in the presence of AcrIIC4. Using previously determined structures of Nme1Cas9 in target DNA bound and unbound states,[19] we generated a model of the AcrIIC4-Cas9 complex that revealed a potential allosteric functional mechanism for this anti-CRISPR protein. This model was supported by mutational data that identified positions in the putative interaction interface for both AcrIIC4 and the Cas9 REC lobe that resulted in loss of anti-CRISPR activity. We propose that the interaction of AcrIIC4 with the REC2 domain inhibits the conformational change required to allow the HNH endonuclease domain to access the target DNA, thus locking the Cas9 in a catalytically inactive state. As the REC lobe is a universal feature of Cas9 proteins and the conformational changes required to bring the HNH and RuvC domains close to the target DNA after it is bound are conserved, this provides a reliable mechanism for anti-CRISPR inhibition.

Since their initial discovery in 2016,[11] 28 unique families of anti-CRISPR proteins that inhibit Cas9-based CRISPR-Cas systems have been described. Mechanisms of inhibition have been determined for nine of these families. The most common mechanism is blocking DNA binding, which is seen in six of the nine families. Inhibition of Cas9 activity after DNA binding has only been observed for two families, AcrIIC1 and AcrIIC3. These two anti-CRISPRs function through similar mechanisms. AcrIIC1 was shown to bind to the active site of the HNH endonuclease domain and trap Cas9 in a catalytically inactive state,[12] while AcrIIC3 was shown to interact with both the HNH and REC2 domains of Cas9.[19,20,30] In this case, two molecules of AcrIIC3 bridge two Cas9 proteins, with each AcrIIC3 monomer forming interactions with the HNH domain of one Cas9 and the REC2 domain of the other, leading to the formation of a Cas9 dimer.[19,20] Thus, AcrIIC3 inhibits cleavage activity both directly through the HNH domain, and indirectly through its interaction with the REC2 domain. The dimerization was originally thought to prevent DNA binding by Cas9,[12,20] but further investigation revealed that AcrIIC3 allows DNA binding and instead inhibits the movement of the HNH domain, preventing it from contacting the DNA cleavage site.[19] Our results show that, like AcrIIC3, AcrIIC4 binding neither depends on nor prevents sgRNA loading and dsDNA binding.[16] Also similar to AcrIIC3, we found that AcrIIC4 interacts with the REC2 domain, and likely inhibits the movement of the HNH domain, thereby blocking DNA cleavage.

The experiments presented here show the AcrIIC4-Cas9 complex binds target DNA but does not cleave it. These results disagree with previously reported results that concluded AcrIIC4 blocks DNA binding in an electrophoretic mobility shift assay.[16] In order to address this discrepancy, we explored DNA binding in the presence of AcrIIC4 using a number of different biophysical and biological techniques, such as electrophoretic mobility shift assay, fluorescence polarization, and *in vivo* transcriptional repression assays. Each of these experiments clearly showed that Cas9 could bind target DNA in the presence of AcrIIC4. There are several possible explanations for the difference observed between the two studies, including the use of different guide RNA and target DNA sequences, as well as differing buffer conditions used in the assays. Similar discrepancies in DNA binding activity were previously observed with anti-CRISPR AcrIIC3, which also interacts with the REC lobe.[12,19,20] In this case, the crystal structure of the DNA-bound AcrIIC3-Cas9 complex showed conclusively that this anti-CRISPR did not block DNA binding,[19] in contrast to the results published in three previous studies.[12,20,30]

CRISPR-Cas9 is a powerful biotechnological tool being developed for many diverse applications in gene editing and gene regulation.[31] For these uses, issues of potential off-target effects need to be overcome. Multiple strategies to improve CRISPR-Cas9 fidelity include directly engineering the enzyme to be more accurate,[32] taking advantage of differing PAM requirements,[18] and using cell-cycle regulated or tissue specific promoters to control CRISPR-Cas9 expression.[33] As many type II anti-CRISPRs, including AcrIIC4,[16] have been shown to successfully inhibit CRISPR-Cas9 activity in mammalian cells, they hold great promise as control switches for CRISPR-Cas9 activity. Continued investigation of the anti-CRISPR landscape will allow the development of unique regulators of CRISPR-Cas9 activity and provide new insight into how anti-CRISPR activity is driving CRISPR-Cas evolutionary diversity.

## Acknowledgments

This work was supported by the Canadian Institutes of Health Research operating grants to K.L.M. (PJT-152918), A.R.D. (FDN-15427), and T.F.M. (PJT-148795).

## MATERIALS AND METHODS

### Phage targeting assays

*E. coli* BB101 cells were transformed with a plasmid expressing the relevant Cas9 protein with an sgRNA containing a spacer that targets phage Mu, and a second plasmid expressing anti-CRISPR. The cells were grown in lysogeny broth (LB) supplemented with 25 μg ml^−1^ chloramphenicol and 50 μg ml^−1^ streptomycin for 2 h at 37 °C. A final concentration of 0.01 mM isopropyl-1-thio-Beta-D-galactopyranoside (IPTG) was added to induce anti-CRISPR expression, and the cells were incubated for 3 h. 200 μL of cells were mixed into 0.7 % LB top agar and poured onto plates of LB supplemented with chloramphenicol, streptomycin, 0.2 % arabinose, and 10 mM MgSO_4_. Serial dilutions of phage Mu were applied to the plates, and the plates were incubated at 37 °C overnight. Assays were performed at n ≥ 3, with representative replicates shown in the figures.

### Nickel affinity co-purification assays

Co-purification experiments with 6His-tagged HpaCas9 constructs and untagged AcrIIC4 were performed as previously described.[13] For the REC2 co-purification experiments with AcrIIC3 and AcrIIC4, a plasmid expressing HpaCas9 REC2 with an N-terminal 6His-tag (pMCSG7) was transformed into *E. coli* BL21 (DE3) cells. An overnight culture of these cells grown at 37 °C was subcultured in fresh LB media supplemented with 100 μg ml^−1^ ampicillin. The culture was grown at 37°C until the OD_600_ reached 0.8-1.0. Then a final concentration of 1 mM IPTG was added to the culture to induce REC2 expression. After overnight incubation at 16°C, cells were harvested and resuspended in the cold binding buffer containing 50 mM HEPES pH 7.0, 300 mM NaCl, 5 % glycerol and 5 mM imidazole. The cell resuspension was lysed by sonication and centrifuged at 17000 rpm for 30 min to remove the cell debris. The supernatant was mixed with Ni-NTA agarose beads (Qiagen) and incubated at 4°C for 30 min. After several washes with the purification buffer containing 50 mM HEPES pH 7.0, 300 mM NaCl and 30 mM imidazole, REC2 was eluted from the Ni-NTA using the elution buffer with 50 mM HEPES pH 7.0, 300 mM NaCl, 5 % glycerol, and 300 mM imidazole. The protein was dialyzed into 50 mM HEPES pH 7.0, 300 mM NaCl, and 5% glycerol. Plasmids (pCDF1-b) expressing untagged AcrIIC3 or AcrIIC4 were individually transformed into *E. coli* BL21 (DE3) cells. The cells were grown as described above in LB media supplemented with 50 μg ml^−1^ streptomycin. After adding 1 mM IPTG and overnight induction at 16 °C, the cells were harvested and resuspended in cold binding buffer. The cells were lysed by sonication and centrifuged at 17000 rpm for 30 min to remove the cell debris. Purified 6His-tagged REC2 was applied to a Ni-NTA column and the anti-CRISPR supernatants were added and the samples were incubated at 4°C for 30 min. Unbound protein was removed through the application of purification buffer to the column and 6His-tagged REC2 and the bound anti-CRISPRs were eluted from the column using elution buffer. The co-eluting proteins were analyzed using SDS-PAGE on a 15 % Tris-Tricine gel visualized by Coomassie staining.

### Purification of AcrIIC4

A plasmid (pHAT4) expressing AcrIIC4 with an N-terminal cleavable 6His-tag was transformed into *E. coli* BL21 (DE3) cells. The cells were grown and AcrIIC4 was purified as described above using buffers with 50 mM Tris pH 7.5, 500 mM NaCl, and 5, 30, or 300 mM imidazole. AcrIIC4 was dialyzed with 6His-tagged TEV protease in 1:100 (w/w) ratio into 20 mM Tris pH 7.5, 250 mM NaCl and 1 mM TCEP followed by another incubation with Ni-NTA at 4°C to remove His-tagged TEV protease. Finally, the protein was injected onto Superdex 200 Increase 10/300 GL size exclusion column (GE Healthcare, USA) equilibrated in Assay buffer (20 mM HEPES pH 7.0, 150 mM NaCl, 5 mM EDTA, 1 mM TCEP, 5 % glycerol) and the purified fractions were collected.

### Purification of Nme1Cas9

A plasmid (pMCSG7) expressing Nme1Cas9 with an N-terminal 6His-tag[34] was transformed into *E. coli* BL21 (DE3) cells. The cells were grown in Terrific broth (TB) and Nme1Cas9 was purified as described above using buffers with 50 mM HEPES pH 7.5, 500 mM NaCl, 5% glycerol, and 5, 30, or 300 mM imidazole. The protein was injected onto Superdex 200 Increase 10/300 GL gel filtration column (GE Healthcare, USA) equilibrated in Assay Buffer (20 mM HEPES pH 7.0, 150 mM NaCl, 5 mM EDTA, 1 mM TCEP, 5 % glycerol).

### HpaCas9 REC and AcrIIC4 cross-linking and mass spectrometry

HpaCas9 REC was purified as described above (Purification of Nme1Cas9). The cross-linking experiment followed the protocol described previously with modifications [XLMS]. A protein concentration of 1 mg mL^−1^ was used and a final concentration of 375 μM of disuccinimidyl suberate (DSS) was added for cross-linking. The cross-linked products were visualized by SDS-PAGE on an 8% polyacrylamide gel and Coomassie staining. The band of interest was excised from the gel and sent for mass spectrometry analysis at the Southern Alberta Mass Spectrometry Facility.

### Luminescence transcriptional repressor assays

A plasmid encoding the J23119 artificial promoter, which drives constitutive expression of the luxCDABEoperon from *Photorhabdus luminescens*,[35] was transformed into *E. coli* BL21 cells together with the pNme1Cas9sgRNA plasmid containing an spacer targeting the constitutive promoter region of J23119[13] and pCDF-1b plasmid expressing the anti-CRISPR proteins. Kanamycin, chloramphenicol and streptomycin resistant transformants were grown in liquid LB medium at 37°C until they reached an optical density at 600 nm (OD_600_) of 0.6. The cells were then further diluted to an optical density of 0.1 in LB, and 100 μl was dispensed into the wells of a 96-well plate. Both luminescence and the OD_600_ were monitored for 24 h using a Synergy H1 reader controlled by Gen5 2.09 software (BioTek Instruments Inc.). The assay was performed at n ≥ 3, with representative replicates shown in the figures.

### Purification of Nme1Cas9-sgRNA RNP

A plasmid expressing Nme1Cas9 with an N-terminal 6His-tag and sgRNA (pMCSG7) was transformed into *E. coli* BL21 (DE3) cells.[11] Nme1Cas9-sgRNA ribonucleoprotein complex was purified as described above (Purification of Nme1Cas9) with the exception that the cells were incubated at 37°C for 3 h after the addition of IPTG. The protein was loaded onto a Superdex 200 Increase 10/300 GL gel filtration column (GE Healthcare, USA) and purified using Assay Buffer or Assay Buffer without EDTA (for use in DNA cleavage assays). The presence of bound sgRNA was assessed by running the protein complex on a 12.5 % polyacrylamide/8 M urea gel and visualizing the nucleic acid using SYBR Gold staining.

### Electrophoretic mobility shift assays

1 μM of Nme1Cas9-sgRNA ribonucleoprotein complex was incubated for 30 min at room temperature with AcrIIC4 or AcrIIC5*_N10_* at concentrations ranging from 0 to 50 μM in Assay Buffer supplemented with 0.01 % Tween-20. 100 nM of fluorescein-labelled target DNA was added and incubated at 37°C for 1 h. The resulting complexes were analysed using a 6% native polyacrylamide gel, and fluorescein-labelled DNA bound to the complex was visualized using a ChemiDoc imager (BioRad).

### DNA cleavage assays

150 nM of Nme1Cas9-sgRNA ribonucleoprotein complex was incubated with 0.1, 1, or 5 μM wild type or mutant AcrIIC4 in cleavage buffer (20 mM HEPES pH 7.5, 150 mM KCl, 5 mM MgCl2, 1 mM DTT, 5% glycerol) at room temperature for 30 min. 300 ng of linearized plasmid DNA containing the target protospacer sequence was added. After a 1 h incubation at 37 °C, the reactions were quenched by adding EDTA and incubating with 1 U proteinase K at 50°C for 10 min. The reaction products were visualized on a 1% agarose gel using SYBR Safe stain (ThermoFisher Scientific).

### Nme1Cas9-sgRNA-DNA-AcrIIC4 size-exclusion chromatography

50 μM of purified Nme1Cas9-sgRNA RNP was incubated with 150 μM of unlabelled target DNA for 30 min at room temperature. 150 μM of AcrIIC4 was added and the mixture was incubated for an additional 30 min. The sample was then injected onto Superdex 200 Increase 10/300 GL gel filtration column (GE Healthcare, USA) equilibrated in Assay buffer. Samples of fractions with peaks at A280 were analyzed for protein components using SDS-PAGE (15% Tris-Tricine gel) and were visualized by Coomassie staining, and for nucleic acids using a 12.5 % polyacrylamide/8 M urea gel visualized by SYBR Gold staining (ThermoFisher Scientific).

### Fluorescence polarization assays

Site-directed mutagenesis was used to engineer a single cysteine residue at the C-terminus of AcrIIC4. This AcrIIC4 variant was purified as described above. AcrIIC4 was then fluorescently labelled using fluorescein-5-maleimide (ThermoFisher Scientific). 4 nM of the labelled AcrIIC4 was incubated with Nme1Cas9 or Nme1Cas9-sgRNA RNP at concentrations ranging from 0 to 4 μM in Assay buffer supplemented with 0.01 % Tween-20 for 30 min at room temperature. Fluorescence polarization values were measured using a TECAN Spark reader.

For the DNA fluorescence polarization assay, 4 nM of fluorescein labelled target DNA was incubated with Nme1Cas9-sgRNA ribonucleoprotein complex at concentrations ranging from 0 to 4 μM in Assay buffer supplemented with 0.01 % Tween-20 for 30 min at room temperature. Fluorescence polarization values were measured using a TECAN Spark reader.

### Crystallization and structure determination of AcrIIC4

Purified 6His-tagged AcrIIC4 was dialyzed overnight at 4°C in 10 mM Tris pH 7.5, 250 mM NaCl and 5 mM βME with 6His-tagged TEV protease in 1:100 (w/w) ratio. The cleaved 6His-tag and 6His-tagged TEV protease were removed by incubation with Ni-NTA beads. The protein was then further purified by size exclusion chromatography using a HiLoad Superdex 75 16/600 gel filtration column (GE Healthcare, USA) equilibrated in 10 mM Tris pH 7.5, 100 mM NaCl.

Crystallization trials of AcrIIC4 were established using sitting-drop vapour diffusion. Purified AcrIIC4 was concentrated to 16 mg/mL, and mixed with the precipitants from MCSG2 and 4, JCSG+, Rigaku Wizard Cryo and Index commercial screens (Hampton Research) in a 1:1 ratio by the Gryphon robot (Art Robbins Instrument). The first crystals were observed in 40% PEG 600, 0.1 M Na_2_HPO_4_/citric acid pH 4.2. The condition was further optimized by altering the polyethylene glycol (PEG) percentage and buffer pH. The crystals formed in 35% PEG 600, 0.1M Na_2_HPO_4_/citric acid pH 4.2 were cryo-protected in the mother liquor with 20% glycerol and flash-frozen in liquid nitrogen. Diffraction data for crystals was collected at the Structural Genomics Consortium (SGC) (Toronto, Canada) using a copper rotating anode X-ray source (lambda = 1.54 Å) and R-AXIS IV++ detector (Rigaku). The phase information of AcrIIC4 was solved by iodide—single-wavelength anomalous diffraction (I-SAD). The data was collected by soaking crystals in the mother liquor supplemented with 20% glycerol and 0.5 M NaI. Diffraction images were processed with mosflm.[36] The model auto-building and automated refinement was performed using Phenix.[37] Manual refinement was performed using Coot.[38]

### Far-UV circular dichroism scans and temperature-induced unfolding experiments

Circular dichroism experiments were performed as described previously[13] with modifications. For the temperature-induced unfolding experiments, the state of protein folding was measured at 207 nm. Assays were performed at n ≥ 3.

### Nme1Cas9 mutants phage targeting assay

In PyMOL, the electrostatics surface map of the Nme1Cas9 structure in complex with sgRNA (PDB: 6JDQ)[19] was generated using the Adaptive Poisson-Boltzmann Solver (APBS) plugin. Asp392, Glu393, Asp394, and Asp418 were identified to in the highly electronegative pocket of the REC2 domain. Point mutations at these positions were introduced in Nme1Cas9 by PCR using oligonucleotides that replace the original amino acid codon with the codon for either alanine or lysine. The introduction of D392A, E393K, D394K, and D418K mutations in Nme1Cas9 were verified by sequencing. The phage targeting assays were conducted as described above.

### Modelling of AcrIIC4 onto Nme1Cas9

In the HADDOCK web server,[28] the Nme1Cas9 structure in complex with sgRNA and target DNA in a seed pairing state (PDB: 6KC7)[19] and the AcrIIC4 structure were used. Nme1Cas9 Asp394 and AcrIIC4 Arg46 were input as the active residues and default parameters were used. The resulting five modelled clusters were analyzed using PyMOL, and the steric clashes were visualized using the show_bumps script written by Thomas Holder.

## ACCESSION NUMBERS

The coordinates and structure factors have been deposited in the Protein Data Bank (PDB ID: <u>**7MSL**</u>).

