## Supplemental Information for "Structural and mechanistic insight into CRISPR-Cas9 inhibition by anti-CRISPR protein AcrIIC4"

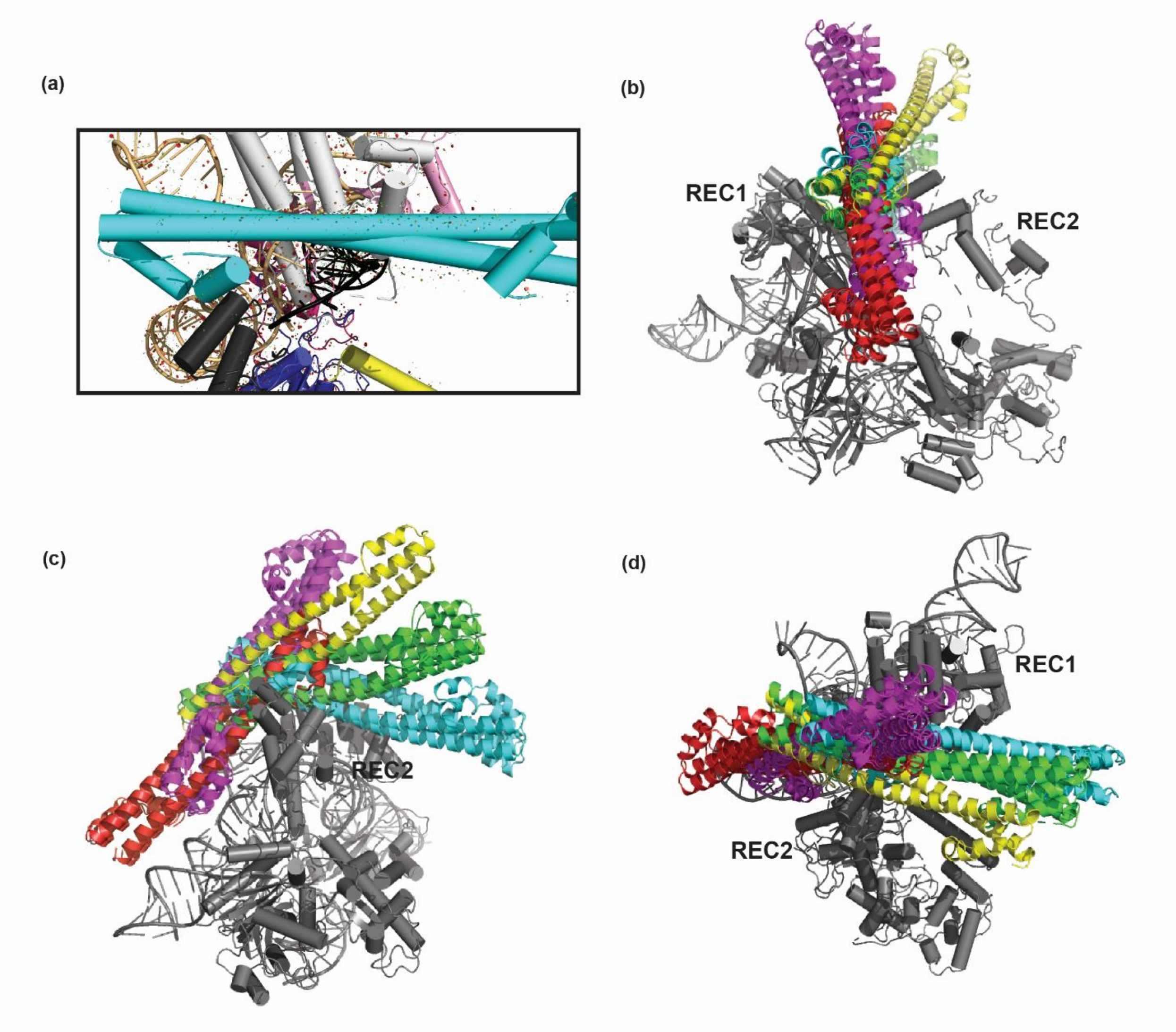


**Supplementary Figure 1**. Model of AcrIIC4 with Nme1Cas9. (a) Close-up of Figure 8(a) with steric clashes between the modelled AcrIIC4 (cyan) and Nme1Cas9 represented with red hexagons. There are minimal steric clashes in the model. (b-d) The five clusters of AcrIIC4 conformations modelled by HADDOCK onto Nme1Cas9 (PDB: 6KC7; gray) are shown in different colors (red, yellow, green, purple, and cyan).

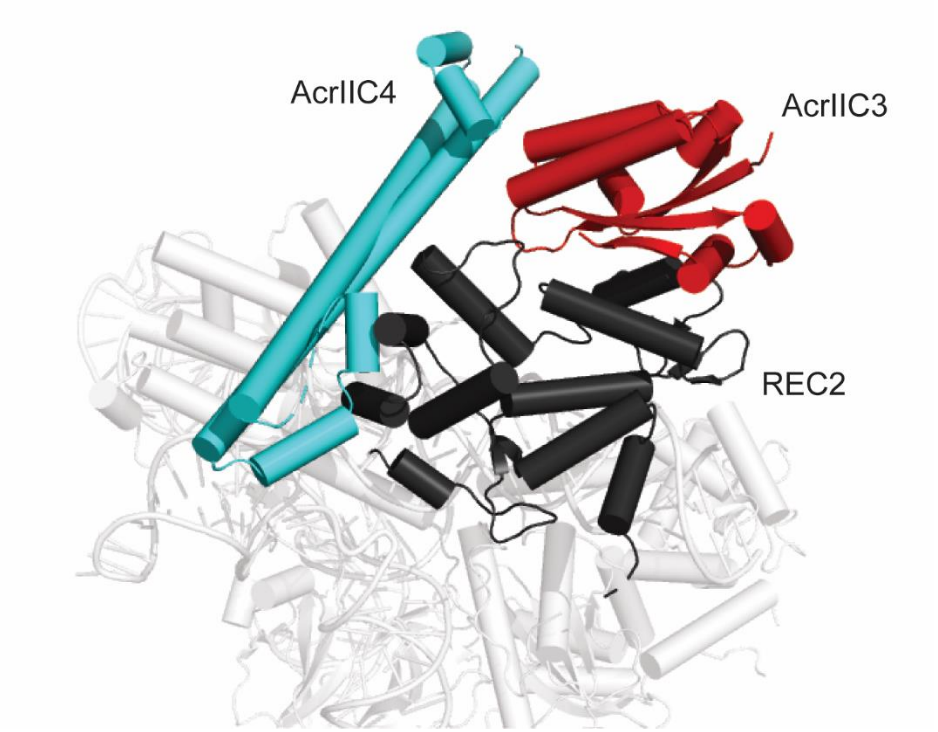


**Supplementary Figure 2.** AcrIIC4 model projected onto the Nme1Cas9 structure bound to AcrIIC3 (PDB: 6JE4). In the crystal structure, two monomers of AcrIIC3 dimerize Nme1Cas9 with each AcrIIC3 monomer binding to the REC2 domain of one Nme1Cas9 and the HNH domain of another. In the figure, only one AcrIIC3 monomer (red) is shown binding to the REC2 domain (black). The model of AcrIIC4 (cyan) is projected onto the structure with no collisions with either the Nme1Cas9 or AcrIIC3.
